# Week-long imaging of cell divisions in the Arabidopsis root meristem

**DOI:** 10.1101/268102

**Authors:** Ramin Rahni, Kenneth D. Birnbaum

**Affiliations:** Center for Genomics and Systems Biology, Department of Biology, New York University, 12 Waverly Place, New York, NY 10003, USA

## Abstract

Characterizing the behaviors of dynamic systems requires capturing them with high temporal and spatial resolution. Owing to its transparency and genetic tractability, the *Arabidopsis thaliana* root lends itself well to live imaging when combined with cell and tissue-specific fluorescent reporters. We developed a novel 4D imaging method that utilizes simple confocal microscopy and readily available components to track cell divisions in the root stem cell niche and surrounding region for up to one week. This new setup allows us to finely analyze meristematic cell division rates that lead to patterning. Using this method, we performed a direct measurement of cell division intervals within and around the root stem cell niche. The results reveal a short, steep gradient of cell division in proximal stem cells, with progressively more rapid cell division rates from QC, to cells in direct contact with the QC (initials), to their immediate daughters, after which division rates appear to become more homogeneous. These results provide a baseline to study how perturbations in signaling could affect cell division patterns in the root meristem.

## Introduction

The transparency and stereotypical growth of the *Arabidopsis thaliana* (Arabidopsis) root has made it an important model to study cell division as it relates to growth, maturation, and asymmetric cell division in a plant meristem (Choe and Lee, 2017). The Arabidopsis primary root tip has a relatively simple but consistent pattern wherein concentric tissue layers all converge on a group of slowly dividing cells—the Quiescent Center (QC) (Fig. 1). Cells in direct contact with the QC, termed initials, behave like stem cells. Initials divide asymmetrically such that one daughter remains in place next to the QC while the other undergoes multiple transit amplifying divisions (Dolan et al., 1993). After a number of transit amplifying divisions, cells elongate, cease dividing, and terminally differentiate.

**Figure 1.**
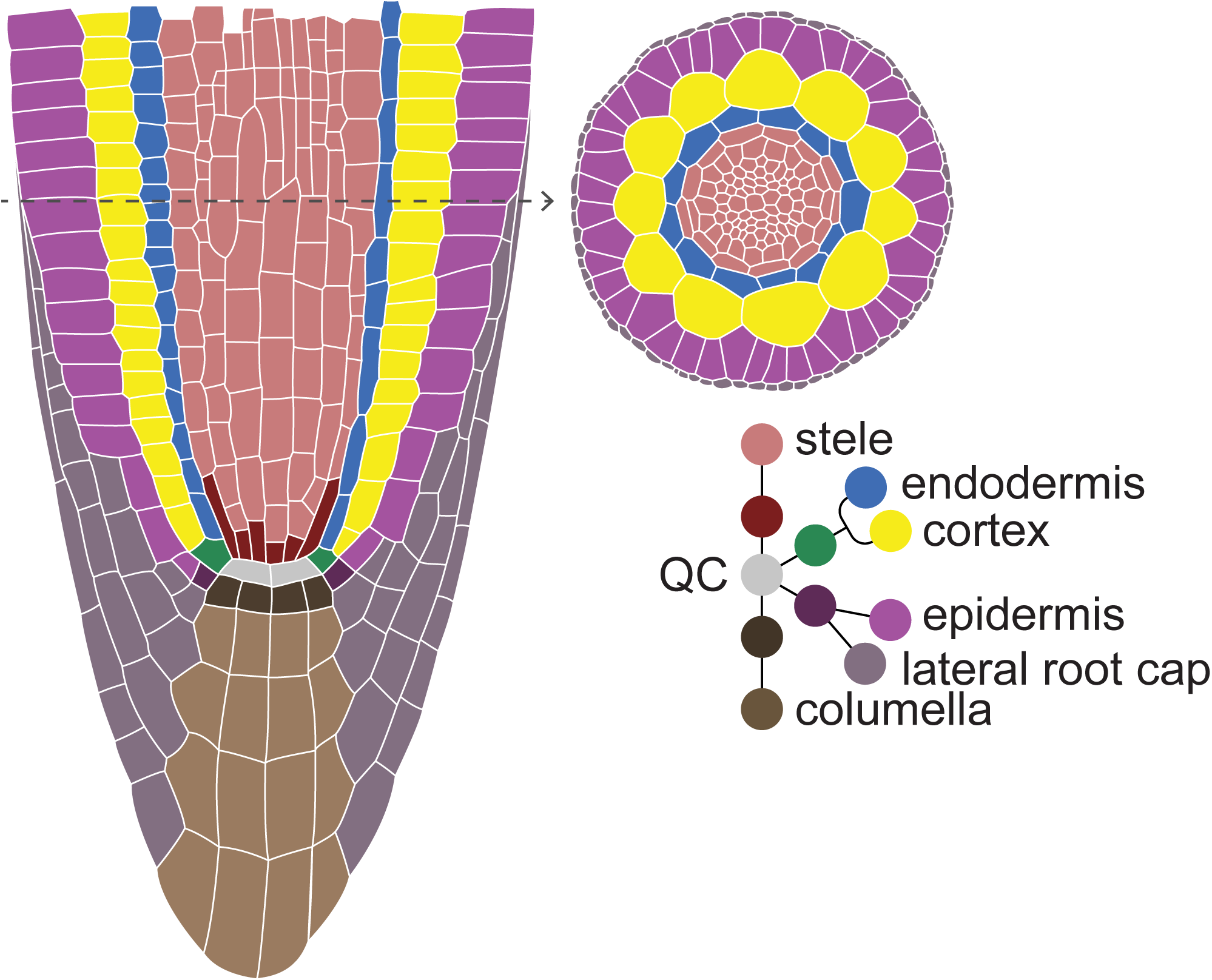
Schematic of a median cross section (left) of the Arabidopsis root apical meristem, showing the tissue-specific stem cells/initials surrounding the QC. Top-right shows a radial view of the root.

While the optical transparency of the root lends itself well to imaging, the requirements for tracking divisions over a long term are challenging. For one, roots need to maintain robust growth while remaining optically accessible for days. In addition, the indeterminate growth of the root means that its position changes dramatically over the time periods needed to study division patterns.

In the Arabidopsi*s* root apical meristem, local division rates vary greatly, apparently reflecting the output of the signaling system of the root. Direct observation of cell divisions for extended periods will shed light on the types of signals that mediate the long-term growth of the plant. Yet, important details about division rates and transitions are not completely understood.

For example, it has been shown that QC cells divide at half the frequency of their surrounding initials, which in turn divide half as often as their transit amplifying daughters (Cruz-Ramírez et al., 2013). However, division rates have classically been estimated from 5-ethynyl-2’-deoxyuridine (EdU) label incorporation or other static methods, with roots imaged at day-long intervals (Cruz-Ramírez et al., 2013) or inferred by imaging several roots for short time periods (Campilho et al., 2006). To better resolve division rates in the meristem under different conditions and backgrounds, a higher temporal resolution is needed, coupled with long-term observation and tracking of individual cells as they divide and grow over several days.

Many groups have developed systems that can track root meristem divisions over extended periods of time. Campilho *et al.* (Campilho et al., 2006) were able to characterize stem cell divisions in the root meristem for over a day on a standard microscope with customized tracking software. They tracked the root mostly in a single central z-plane at each time point, with the longest continuous experiment spanning 28 hours. Another approach has been to use specialized growth systems with, in some cases, specialized microscopes. For example, we have previously developed a custom light sheet microscope with customized tracking for imaging multiple z-stacks over 40 hours (Sena et al., 2011). A more recent approach—MAGIC for the ZEISS Lightsheet (de Luis Balaguer et al., 2016)—imaged roots up to 48 hours in a system that allows for efficient multiplexing of up to 12 seedlings. Similarly, von Wangenheim *et al.*, (2016) imaged lateral roots for up to 64 hours in another light sheet system, also using liquid perfusion of media (Wangenheim et al., 2016). Lastly, the RootArray involves germinating seeds in a specialized device that could be mounted on an upright confocal microscope, tracking roots for 54 hours (Busch et al., 2012).

While these devices were designed for specific applications like parallel imaging of many specimens, we sought to develop a system in which we could track cell divisions and fluorescent markers for up to one week to directly visualize division rates in and around the stem cell niche, which can be exceedingly slow. For our purposes, on-stage perfusion devices limited experiments by requiring highly specialized setups, preventing access to the specimen, or reducing viability of the tissue over time. Thus, we sought a simple, on-stage growth system that utilized solid growth media.

In developing a long-term imaging system that addressed the above-mentioned issues, we realized that a standard confocal microscope and tracking software could be used for rapid interval imaging over many days. We also aimed to keep materials and devices low-tech to develop a system that could easily be adapted in whole or in part by other labs for long-term imaging of the root meristem.

We describe here a novel system that uses readily available components and confocal microscopes for long-term, 4D imaging of roots with high temporal and spatial resolution. By imaging confocal z-stacks of half or more of the root’s volume at roughly 13-minute intervals, every mitotic event in most cell lineages in the field of view can be captured for up to seven days, without significant photobleaching or toxicity.

In addition, we mapped fine-scale patterns of cell division in the meristem to demonstrate the capability of the system and to provide a baseline to address models of cell division control. For example, we have previously proposed a model in which cells may exist along a gradient of division rate from QC to differentiated daughters (Rahni et al., 2016). Using this newly developed live imaging technique, we mapped mitotic rates in each cell position within the stem cell niche and into the early meristem. We show that initials in direct contact with the QC do exhibit a distinctly slower rate of division compared to their daughters, which, in turn, showed a slightly lower rate of division than *their* daughters, forming a short, steep gradient over four cells.

## Results

Our goal was to develop a simple long-term imaging method that takes advantage of commonly available microscopes, materials, and tracking software. To do so, we used a well-known commercial platform (an inverted Leica SPE confocal), but the methods should be adaptable to other systems. The techniques described here meet several important requirements for long-term growth with standard equipment: (i) plants must survive and remain healthy on a microscope slide for up to seven days and growth needs to be restricted to the X- and Y-axes to avoid the need for add-on devices and to allow the use of built-in automated stage controls; (ii) the growing root requires reliable, automated tracking as the tip rapidly moves out of the field of view; (iii) photo-bleaching and -toxicity must be minimized in order to achieve high temporal resolution while applying laser excitation frequently enough to capture relatively short-lived events, such as mitoses, which serve as visual landmarks for keeping track of divisions in individual cell lineages.

### Plant Growth and Survival

To accommodate plant growth and survival, roots were placed between a full-length (25 × 75 mm) rectangular glass #1 coverslip and a block of high-density agar (Fig. 2, A-D; Supplemental Fig. S1). In this set up, plants are restricted to the X- and Y-axes both by the weight of the agar slab and by the apparent inability of roots to penetrate the agar block above. Leaves and hypocotyls are positioned outside the agar and moisture is maintained in the microenvironment by covering the agar and leaves with a small plastic lid (Fig. 2, E **and** F; Supplemental Fig.S1B). As an extra measure, medical micropore tape is used to seal the points of contact between the lid and coverslip (Fig. 2G; Supplemental Fig. S1C).

**Figure 2.**
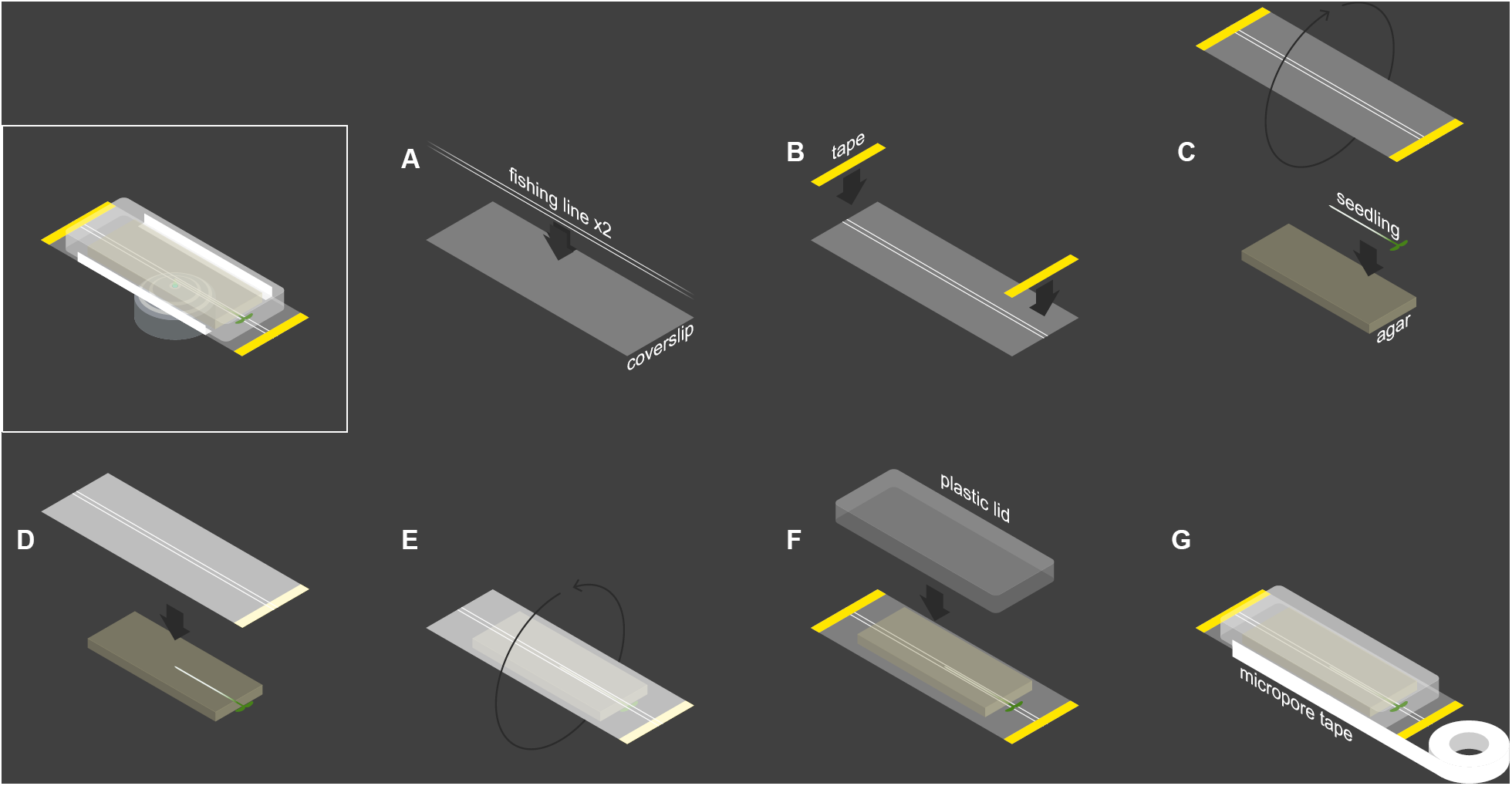
Assembling the growth chamber. A, two strands of fishing line are placed against a full-length coverslip, approximately 1 mm apart. B, the fishing line is affixed to the coverslip with thin strips of laboratory tape. C, seedling is laid against agar block while coverslip is rotated such that the fishing line side faces the seedling. D, coverslip is laid against seedling/agar, positioning the root inside the 1 mm gap between fishing line strands. E, seedling, agar, and coverslip combination are rotated such that the agar side is facing upward. F, a plastid lid is placed over the seedling and agar block to prevent them from drying out. G, the points of contact between lid and coverslip are sealed with micropore tape to further ensure moisture retention and prevent contamination

Agar slabs were prepared by pipetting autoclaved media preparations into square, 12 cm × 12 cm plates and allowing them to solidify before cutting out roughly 2 cm × 4 cm rectangular slabs using a sterile blade. The basal surface of the agar that had been in contact with the bottom of the square plate was placed in contact with the root and cover slip, as this side is freer of heterogeneities compared to the air-exposed surface. This smoothness helps to further minimize root drift in the *z* dimension. Bubbles and gaps between the glass-agar interface are smoothed out by sweeping the agar with a pair of broad tweezers, applying gentle sweeping pressure.

We initially tested different preparations of standard 1/2 MS growth media with various sizes and densities of agar slabs to find an optimal volume and concentration. At 3% or more agar by volume, the slab quickly dried out and roots grown on this preparation showed a significant increase in root hairs, suggesting stress (data not shown). On the other hand, preparations with less than 2% agar by volume lost their firm consistency over time, making them more prone to penetration by growing roots, and more difficult to manipulate when placing roots into the agar/coverslip interface. Larger volumes tended to work best, as the agar—left at room temperature on the microscope stage—slowly deforms, and this deformation is likely accelerated by the persistent higher energy laser light. The largest block that could be contained within our plastic lids was roughly 4 mm in height, which was generated by cutting the block as described above from a 12 cm × 12 cm plate poured with 40 ml of media. We thereafter used 40 ml of media containing 2% agar by volume, which provided an environment on top of the stage that consistently allowed roots to grow without showing visible signs of stress.

### Automatic Tracking of the Growing Root Tip

Given that we wanted to follow cell divisions for a minimum of several days, we required an automated tracking system to follow the root tip as it grew. We sought to adapt a standard tracking system—Leica LAS AF’s MatrixScreener module, which, like most commercial software trackers, is largely optimized for animal cells that migrate limited distances. This contrasts with growing root tips that move out of field considerably faster due to cell expansion. Although primarily designed to follow a single object-of-interest in a chamber of a multi-well plate, we adapted the MatrixScreener module to enable tracking of fast-moving root tips as follows.

During each tracking “job” (see Methods), the MatrixScreener software searches for the brightest plane in the field of view and centers this plane before acquisition. A root tip-specific fluorescent reporter generates a “blob” that can serve as a target to follow. An important feature of the tracking software was on-the-fly drift correction, which was accomplished by setting the plane with highest fluorescence as a reference plane. Note that, while a columella or cap reporter (e.g. *PET111*) can provide a discrete entity to track, its expression spans a rather large volume that presents several bright planes that the autofocus can detect. This can lead to cycling between these planes during drift correction, causing “lurching” of the captured image, as well as unreliable stack registration. Smoother tracking can be achieved with a smaller fluorescent region that provides a reliable visual anchor for the tracking and drift correction modules. We used the QC-specific reporter *pWOX5::GFP(ER)* (Blilou et al., 2005), which strongly labels a small cluster of cells, allowing for stable tracking and a small margin of error during drift correction. With a 20x objective at 2.5x zoom, this setup permits a field of view typically spanning 2/3^rds^ the cells of the meristem, about 15 cells from the QC.

Roots can grow in any orientation and have a natural tendency to twist as they grow, causing cells to rotate into the positions farthest from the objective and most difficult to view (see below). In addition, a longer, horizontally growing root can coil if the tip grows back in on itself (Supplemental Fig. S2; Supplemental Movie S1). We observed that the twisting action was minimized when the root grew alongside a straight object, which it naturally tends to “hug.” Thus, we stabilized the growth vector of the root using a pair of on-slide “bowling bumpers” made of commercial fishing line and spaced roughly 1 mm apart (Fig. 2A). These guides minimized twisting and eliminated coiling (Supplemental Fig. S3; Supplemental Movie S2), greatly enhancing the ability to track the same lineage of cells over many days. While other types of fishing line may be used, we note that higher diameter lines seem to create more of a pocket for moisture along its edges, which can cause optical aberrations and dark patches, and roots can occasionally grow over lines with smaller diameters, exiting the guide alley.

### Photobleaching and Phototoxicity

One challenge in long-term imaging is avoiding photo-bleaching and -toxicity, especially in this case, where short intervals between exposures are needed to track cell divisions. These effects were mitigated by using low laser power (20% or less) and limiting acquisition loops to an interval of roughly ten minutes between the end of one loop and the start of the next to allow sufficient recovery time. These settings allowed us to reliably track individual cells (using the *35S::H2B-mRFP1* reporter (Federici et al., 2012)) within about 2/3^rds^ of the meristem closest to the objective. Acquisition intervals were specified by observing the runtime of the loop for that given set of imaging settings and adding 10 minutes to this time. For example, sequentially imaging stacks of both a GFP and RFP reporter may take 3 minutes, corresponding to a 13-minute total imaging interval. Importantly, this interval is short enough to reliably observe all mitotic events in any visible cell over the entire time course.

Experiments using multiple laser lines will therefore require larger refractory windows, although we have used periodic exposure to other laser lines to observe cell fate markers, for example, during time lapse imaging. In general, windows longer than 15 minutes may not be short enough to catch landmark events with short lifetimes (e.g. mitoses) and are less reliable to follow any particular cell over several time points. A high scan rate (600 Hz) carried out in bi-directional mode (see Methods) helps further expedite the acquisition process, and each channel is averaged a total of three times to clean up background noise. Protein fusions with a higher turnover rate (such as the membrane-localized aquaporin *pLti6b:GFP* (Federici et al., 2012)) were less prone to bleaching effects than fusions to more stable proteins (such as the Histone 2B reporter *35S::H2B-mRFP1* (Federici et al., 2012)) (data not shown).

To test photo-bleaching and -toxicity in the system, we measured overall division rates in the meristem as an indicator of meristem viability and fluorescence longevity from an endogenous promoter in early and late time points under a roughly 13-minute exposure regime. To assess photobleaching, we calculated Corrected Total Cell Fluorescence for *pWOX5::GFP(ER)* expression at a median z-section for each of the four films, sampling six time points from the start of image acquisition to 60 hours later (a time point common to all four films). We observed some photobleaching during the first 24 hours, but fluorescence stabilized and gradually recovered over the time course (Supplemental Fig. S4). To assess phototoxicity, we counted cells that were either about to undergo mitosis, were in mid-mitosis, or had just finished dividing at a median section roughly every two hours from film start to 65 hours later. The frequency of mitoses drops slightly from 5 per time point to 4 in the early time points as plants appear to adjust to the new growing condition, but then division rates stabilized and remained at a constant level (Supplemental Fig. S5). These results indicate that, in the imaging system, growth remained stable and, while an endogenous marker showed some early bleaching, it did recover and could be imaged for long periods.

### Characterizing Divisions in the Stem Cell Niche and Transit Amplifying Zone

Division rates in the Arabidopsis meristem have been shown to progressively increase from the QC to initials to transit amplifying cells. For example, using S-phase incorporation methods, relative division rates were approximated to be N, 2N, and 4N, for the QC, initials, and transit amplifying (TA) cells, respectively (Cruz-Ramírez et al., 2013), and division rates in the TA zone were inferred to be 17 hours (Cools et al., 2010; Hayashi et al., 2013). This relatively lengthy duration of the cell cycle, and overall quiescence of the QC in particular, have made direct measurements difficult. In addition, it has not been clear if differences in division rate are truly discrete, or, if some kind of gradient might exist. This distinction has important implications about the nature of the signals that control division rates. Moreover, these broad characterizations tend to average data from several tissues, potentially missing tissue-specific patterns.

To more finely characterize the mitotic behavior of cells in the root meristem, we used our live imaging method to record cell divisions in the meristem over several days, including up to one week. This long imaging window allowed us to follow relatively slow-cycling cells as they went through sequential divisions, giving direct measurements of mitotic rates.

The combined results from four separate time lapse experiments (spanning 72h, 96h, 116h, and 168h; Supplemental Table S1) are summarized in Figure 3. Each filled circle ● represents an instance where we observed two sequential, or “bookended,” divisions of the same cell—“hours” corresponds to observed division interval. Empty circles ○ correspond to cells that divided only once over the course of the film but could be observed for a long period of time in the movie either before or after their first division. For these empty circle data points, “hours” represents the time between the observed division and the beginning or end of the film. We included these data in the first-pass analysis for initials and QC since it provides more data on the *minimal* interval between divisions of the initial. We refer to these intervals as division rates.

**Figure 3.**
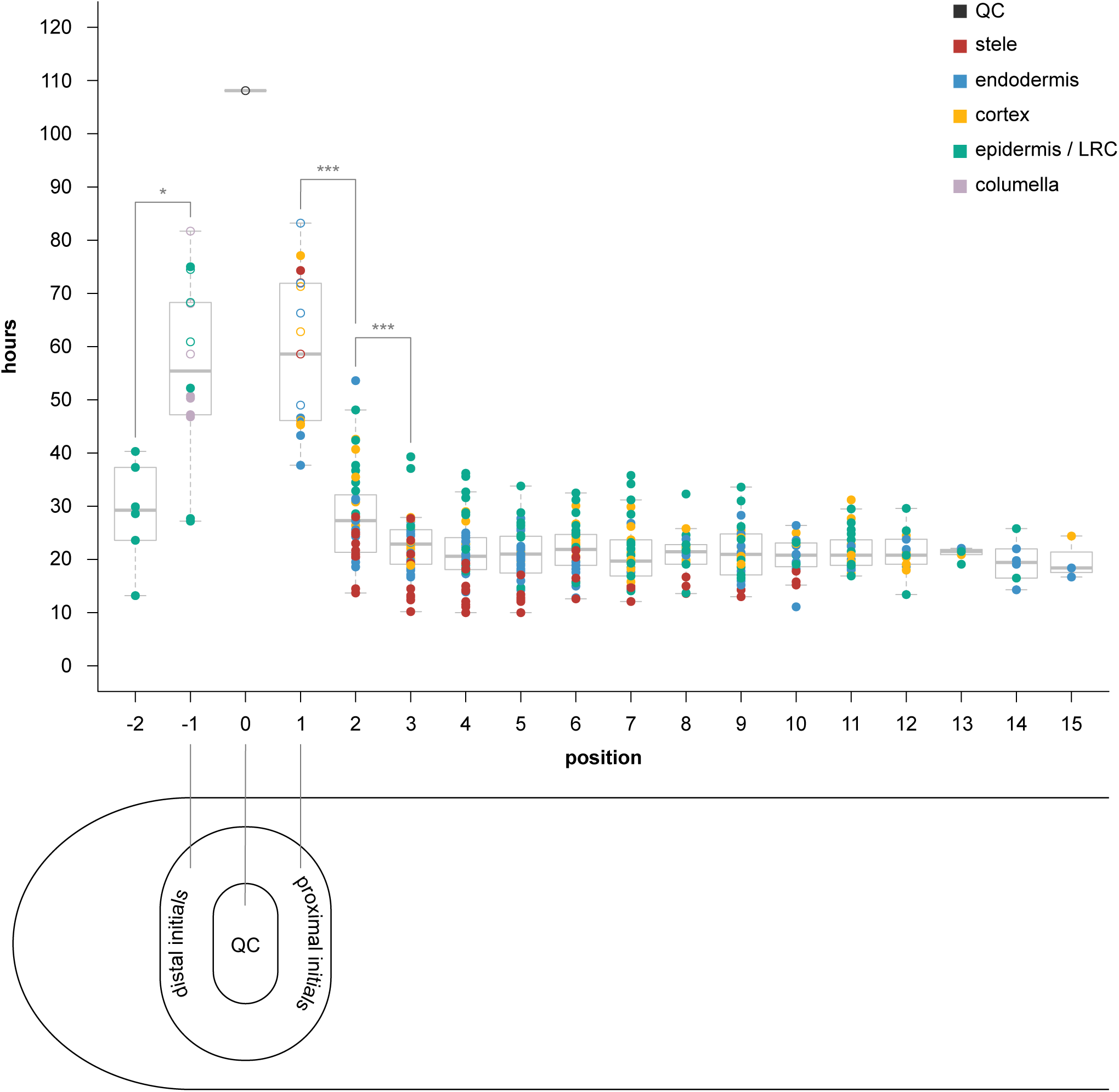
Results of time-lapse experiments, showing hours between divisions by cellular position. Cells are color-coded by tissue type (legend in upper right-hand corner). Closed circles represent two “bookended” observed divisions for the same cell. Open circles denote a cell with only one observed division throughout the course of the film, with “hours” representing the time from that division until the beginning or end of the film (whichever happened to be longer). QC is denoted position 0. Proximal initials (stele, endodermis, cortex) are position 1. Distal initials (columella, epidermis/lateral root cap) are position −1. Positions 2 and -2 are the immediate daughters proximal and distal initials, respectively. Four biological replicates have been used. Stars represent statistical significance (*: p<0.01, ***: p<0.0001).

The QC position is labeled “0”. Cells in direct contact with the QC—initials—are labeled “1” or “-1”, for proximal or distal stem cells, respectively. The neighboring daughter cells of initials are labeled “2” or “-2”, and so forth. Note that we did not observe a distinct cortex/endodermis initial (CEI) in any of the QC-adjacent cells we were able to track for the entire duration of the four films. Therefore, the cortex and endodermis cells directly adjacent to the QC are labeled separately with a designation of “1”—the initial position. Additionally, while the epidermal/lateral root cap (LRC) initial can alternate between anticlinal and periclinal divisions to generate these two tissue types (Wenzel and Rost, 2001), we only observed periclinal (LRC-generating) divisions of this cell type in our films.

A two-way Analysis of Variance (ANOVA) revealed a strong effect on division rate by both position (p < 0.0001) and tissue (p < 0.0001), but not by their interaction (p = 0.86) (Supplemental Fig. S6A), which indicates that position-by-position trends in division rates are similar among tissues despite differences in the magnitude of division rates.

Focusing on the proximal meristem data (positions 1-15), we found that proximal initials (position 1) had a median division interval of 58.6 hours (mean: 58.02) (example: Supplemental Movie S3), while the immediate daughters of proximal initials (position 2) had a significantly faster division rate (ANOVA, Tukey Test, p < 0.0001, Supplemental Fig. S6B), with a median of 27.3 hours (mean: 28.3), showing a sharp increase in division rates after being displaced only one cell length away from the QC (Fig. 3). Interestingly, cells at position 2 showed a smaller but significant change in division rate compared to their proximal neighbor (position 3)—27.3 vs 22.9 hours (means: 28.3 vs 22.27, respectively) (Tukey Test, p < 0.0001)—suggesting another one-cell length speeding up of division rates, albeit less dramatic than the preceding interval (Fig. 3).

Comparing distal initials (position −1) with their immediate daughters (position -2), we also found a significant difference in division rate (t-test, p < 0.01) (Fig. 3).

Note that these calculations include the data points represented by hollow circles indicating minimal division intervals in Fig. 3. If only division intervals with bounded mitotic events (filled circles) are included, the median rate for positions -1 and 1 combined is faster at 47 hours (mean – 50.1; data not shown). Nonetheless, the difference between initials (positions -1 and 1 combined) and the proximal cells (2-15 combined) is still statistically significant (Tukey Test, p < 0.0001), as is the difference between positions 1 and 2 (Tukey Test, p < 0.0001). The data between positions 2 and 3 is unchanged, so still represents a significant change in division rates.

The difference between positions 3 and 4 was not statistically significant (Fig. 3; Supplemental Fig. S6B). Indeed, median division rates in the proximal zone after position 3 appear constant, with no significant differences among them using the Tukey Test (Fig. 3, Supplemental Fig. S6B). In several tissues, there does appear to be a very subtle trend toward more rapid cell division rates up to position 5 (Supplemental Fig. S7). Thus, division rates were either flat or speeding up at a much more gradual rate after position 3.

The transit amplifying population (positions 2-15) pooled together had a median of 21.5 hours (mean: 22.15) between divisions (example: Supplemental Movie S4). This reflects a 2.7-fold difference in division rate between initials (positions -1 and 1 combined) and transit amplifying cells (2-15 combined).

The data also revealed wide variation in division rates at all positions. While the median time between divisions in the TA zone (positions 2-15) was 21.5 hours, cells divided as rapidly as every 10 hours and as slowly as every 53.6 hours within the TA zone (Fig. 3; Fig. 4). The last five or so positions in the meristem show a smaller range of values (Fig. 3) but this is likely due to the fact that many fewer sequential divisions could be observed as cells exited the frame.

**Figure 4.**
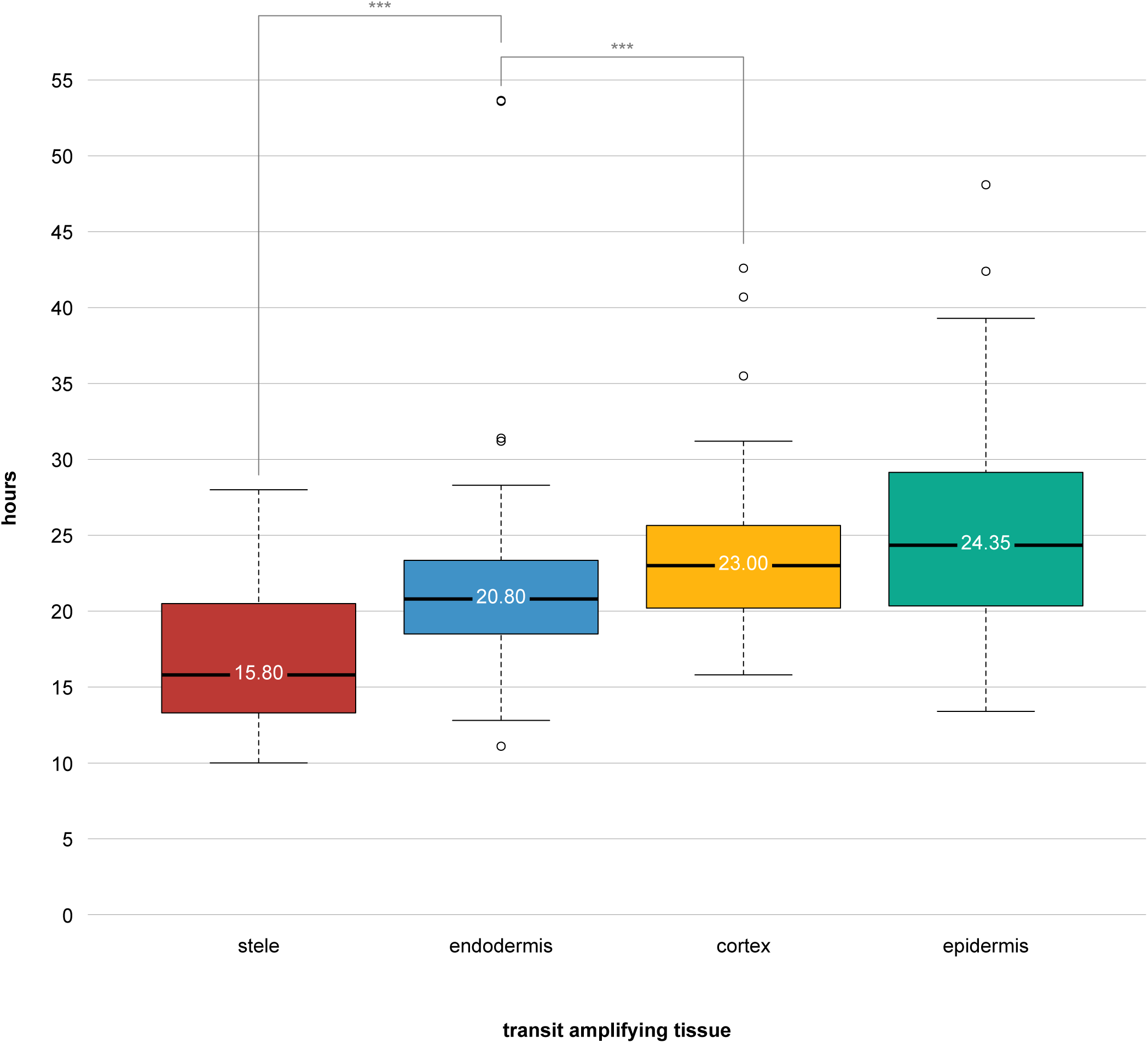
Results of division times in the proximal meristem (positions 2-15 in Figure 3), sorted by tissue type. The value displayed on each box plot represents the median. Stars represent statistical significance (***: p<0.0001).

Much of the variation was due to a radial gradient of division rates. Focusing on the TA zone data (positions 2-15), we found significant differences in tissue-specific division rates (ANOVA, p < 0.0001, Supplemental Fig. S6C), with the fastest divisions in the stele (median: 15.80 hours, mean: 16.97 hours), followed by the ground tissue (endodermis—median: 20.80 hours, mean: 21.30; cortex—median: 23.00 hours, mean: 23.46 hours) and then the slowest rates in the epidermis (median: 24.35 hours, mean: 25.25 hours) (Fig. 4). A Tukey Test on these TA cells revealed all tissues having significantly different division rates, except cortex and epidermis, which were not significantly different from one another (Fig. 4; Supplemental Fig. S6C). Thus, there appears to be both a proximo-distal and a radial gradient of divisions that are spatially separated.

Of the QC cells that we could follow for the entire duration of the film, only a single QC division was observed (Fig. 3; Supplemental Movie S5). The four films combined span a total 452 hours of observations, suggesting a mitotic frequency potentially greater than 7 days in our growth system. This is consistent with previously reported pulse-chase experiments showing F-*ara*-EdU label retention in the QC up to 4 days after chase (Cruz-Ramírez et al., 2013). The single observed QC division generated a daughter cell in the cortex/endodermal initial (CEI) position (Supplemental Movie S5). Interestingly, *WOX5* expression appeared diminished in this CEI-positioned cell and its QC parent as compared to the neighboring QC cells.

Overall, the high-resolution analysis of division rates indicated a short gradient over four cell lengths characterized by a dramatic increase in cell division rates from QC to initial to immediate daughter and then a much less dramatic increase in rate to second daughter. Thereafter, division rates within the 15 or so cells we could observe in the meristem appeared homogenous, or nearly so.

## Discussion

We have developed a relatively straightforward method for live imaging Arabidopsis roots for several days, using readily available materials and a commonly available microscope. The protocol—consisting of a specific growth environment, excitation settings, and tracking parameters, along with transgenic lines with a nuclear reporter—allows for the study of cell division behavior and dynamic lineage analysis over multiple days. Moreover, it can be utilized with plants of different genetic backgrounds or media conditions to study genetic and environmental effects on the meristem. The automated tracking component should allow long-term imaging, where users in shared facilities can set up experiments overnight or over weekends.

Plant health does not seem to be dramatically affected by the growth conditions and laser exposure. Indeed, long term fluorescent readouts from native promoters are possible in this system, but some care needs to be taken for quantitative readouts of native reporters because of the potential for early partial bleaching followed by stabilization.

While the relative quiescence of the QC and initials was known, the division patterns within and nearby the stem cell niche had not been fully characterized, particularly among the proximal daughters of the initials. Using our method, we found a sharp transition between the division rates of the initials and their immediate daughters and another increase in division rates in the next set of daughter cells (Fig. 3). This sharp but steep gradient from QC to position 3 is followed by more uniform division rates, although there are overall tissue-specific differences in division rates (Supplemental Fig. S7). In addition to the proximal gradient, we found that initials divide nearly 3-fold more slowly than TA cells, which is slightly more pronounced than previously estimated rates (Cruz-Ramírez et al., 2013), although growth conditions in the two experiments were different. In addition, the median time between divisions in the TA zone was 21.5 hours, slightly slower than the previously estimated 17 hour duration of the cell cycle (Cools et al., 2010; Hayashi et al., 2013).

The direct measurement of division rates in individual cells also permitted an assessment of the range and variance of cell cycle durations in the meristem. These data showed that cells exhibit a five-fold range in division rates among TA cells (Fig. 3 and Fig. 4), much of which was explained by tissue- and cell type-specific rates of division, with a gradient of increasingly slower division rates moving outward in the radial dimension. These results show an inverse relationship between cell size and division rates among transit amplifying cells (Fig. 4), which is intuitive since the overall expansion rate of inner and outer tissues of the root at a given point in the maturation zone must be equal (De Vos et al., 2014).

In addition, despite being the tissue with the most rapid TA division rates on average (Fig. 4), the stele seems to have relatively slow cycling initials (Fig. 3). Low-level *WOX5* transcription has been reported in the stele initials (Pi et al., 2015) and so this discrepancy may reflect some basal *WOX5* activity in these cell types, which would be consistent with its role in maintaining relative quiescence via repression of *CYCD3;3/CYD1;1* (Forzani et al., 2014).

Of the QC cells we were able to follow for the duration of each film, only one divided. With the longest film being 1 week long (168 hours), this suggests that QC cells have a mitotic frequency that is at least nearly 3x slower than initials and nearly 8x slower than TA cells. We note, however, that QC division intervals may even be much longer, since the minimal estimate was dictated by our observable window—that is, seven days. We also note that different environmental conditions may affect both overall division rates and relative division frequencies within the meristem. Our growth system would allow different media conditions to be tested to determine how they affect division patterns.

The one observed QC division (Supplemental Movie S3) gave rise to a cell in the CEI position. The QC can act as a reservoir to replace lost or damaged stem cells (Heyman et al., 2013), although its primary contributions appear to be the columella (Cruz-Ramírez et al., 2013). We did not observe any cells in the CEI position in any of our tracked cortex or endodermis lineages. It remains possible that this cell type is transient, and only occasionally replenished by QC divisions. Further analysis is needed to clarify these two issues.

Direct contact with the QC is thought to maintain initials in a stem-like state. WOX5 protein was recently shown to be a mobile transcription factor that moves one cell layer over into the distal columella stem cells to repress their differentiation and ostensibly maintain their stem-like state (Pi et al., 2015). Still, much less is known about what controls the properties of proximal stem cells.

The distinct division rate of the proximal initials that we found is consistent with an as-yet-unidentified QC-sourced signal influencing division rates in these cells. However, the source of the signal is not clear, as our results do not rule out that sources other than the QC could control proximal division rates (Rahni et al., 2016).

It remains to be tested, for example, if mutants perturbed in QC identity but displaying near-wild type growth rates (Sebastian et al., 2015) possess wild type division patterns in the proximal initials and their daughters. Or, if a general block of signaling from the QC (Liu et al., 2017) affects proximal initials and the short gradient of division we characterized here.

Importantly, the distinct division rates of the proximal initials compared to their daughter cells and the QC constitutes a specific property of these initials. We note that this behavior could be used as a marker to assess the effect of perturbations on proximal stem cells and how their division patterns are coupled or uncoupled from that of their neighbors.

The fine-scale properties of division rates in the meristem will help dissect the signals and sources that control division rates and longevity—a central feature of plant growth. Fine-scale characterization of mutants and other perturbations will help answer whether there is a localized population of cells (e.g. QC) that controls division activity around the meristem, or whether multiple sources of signals fine-tune divisions in different regions. Addressing these questions will ultimately help us understand how plants, as sessile organisms, can grow for such long periods of time. Overall, these data show the potential of a convenient, long-term imaging system that can directly measure division rates to analyze important properties of meristem growth and function.

## Conclusions

We have developed a novel but simple method for live imaging plant roots over several days. Using only readily available components, it is accessible to a broad range of plant researchers. By tracking cell divisions, we have quantified the mitotic rate of various tissues and positions within the Arabidopsis root apical meristem. We show the cell-by-cell resolution of a sharp gradient of division rate from QC to second stem cell daughters. Additionally, our data show a radial gradient in division rates in the TA zone, from fast in the stele to slower in the epidermis. This method may be combined with various genetic and pharmacological perturbations to further dissect the division behaviors in the stem cell niche and surrounding region.

## Materials and Methods

### Plant Growth and Materials

Seeds of Arabidopsis plants carrying the transgenes *35S::H2B-mRFP1* (Federici et al., 2012), and *pWOX5::GFP(ER)* (Blilou et al., 2005) (both in Col-0 background) were stratified at 4°C for 2 days, sterilized, placed on agar plates containing 1/2 Murashige and Skoog salts (Sigma M5524), 0.5% sucrose, and grown vertically in chambers set to 23°C and a 16h light / 8h dark cycle (80-90 µmol m^-2^ s^-1^). Seedlings were transferred to microscopy chambers at 4 days post germination, except for the seven-day long imaging experiment in which case they were transferred 2 days post germination. Roots were allowed to acclimate to the chambers, allowing time to “hug” one of the fishing line guides; while several hours can be sufficient, overnight acclimation in the growth chamber gives the most reliable results.

The small plastic lid used in these experiments was a cover from the Lab-Tek^®^ Chamber Slide^™^ System 177410. The medical micropore tape used was 3M Micropore^™^ 1.25 cm REF 1530-0. The “bowling bumpers” were made with GOTURE 500M 4LB 0.10mm 1.80KG #0.4 monofilament fishing line.

### Microscopy

Arabidopsis roots with fluorescently labeled nuclei and QC were imaged every roughly 13 minutes over the course of 3-7 days on a Leica SPE inverted confocal microscope. All experiments used an air objective (20x/0.7) with 2.5x zoom within the LAS AF software. The time intervals between acquisitions and the laser power and gain used per channel are:

<u>72 hour film</u>: 13.17 minute intervals; 15% 488 nm laser power – 900 V gain; 20% 561 nm laser power – 900 V gain

<u>96 hour film</u>: 13 minute intervals; 15% 488 nm laser power – 905 V gain; 20% 561 nm laser power – 905 V gain

<u>116 hour film</u>: 13 minute intervals; 15% 488 nm laser power – 905 V gain; 20% 561 nm laser power – 905 V gain

<u>168 hour film</u>: 13 minute intervals; 20% 488 nm laser power – 900 V gain; 20% 561 nm laser power – 915 V gain

### Image Processing

Confocal stacks spanning over half the root were acquired at each time point, and stacks were registered using the Correct 3D Drift plugin in FIJI (Parslow et al., 2014). Each cell lineage was followed by eye, beginning at time 0 (film start) and until either the end of the film or until they were displaced out of frame. Lineages were each followed at least three times, making sure to follow them both forward in time and reverse. To keep note of individual lineages, trees were constructed using the Newick convention and visualized using PhyD3 (Kreft et al., 2017).

All main figures were created in Adobe Illustrator CC 2018 with Figure 3 and Figure 4 first being exported from R as SVG files and then assembled and overlaid in Illustrator.

### Video Processing

Maximum projection films were generated in Leica LAS AF using the built-in Processing tools. After image registration in FIJI, all films were exported as .avi files and brought into Adobe After Effects CC 2018 to add text, arrows, time stamps, and other annotations. The green channel for all films was recolored to cyan in After Effects to make accessible for those with red-green color vision deficiencies (Wong, 2011) using the settings below:

#### Change to Color

From: #00FF00

To: #00FFFF

Change: Hue

Change By: Setting To Color

Hue: 33.0%

Lightness: 100.0%

Saturation: 100.0%

Softness: 100.0%

Supplementary Movie S5 required zooming in on an already low resolution film and so further processing was carried out after zooming to make the events easier to distinguish by eye:

#### Color Balance

Highlight Red Balance: 10.0

(all else 0.0)

#### Sharpen

Sharpen Amount: 25

#### Brightness & Contrast

Brightness: 0

Contrast: 25

### Statistical Analysis

Statistical analyses were carried out in R, with the ANOVAs and Tukey Tests done in MVApp (Julkowska et al., 2018), which can account for different number of levels within each factor. The data used for all analyses is available in Supplemental Table S1.

Results from ANOVA and Tukey Test for all positions (−2, −1 & 1-15) are listed in Supplemental Fig. S6A and correspond to rows 2-21, and 23-388 in Supplemental Table S1.

Results from ANOVA and Tukey Test for proximal domain (positions 1-15) are listed in Supplemental Fig. S6B and correspond to rows 23-388 in Supplemental Table S1.

Results from ANOVA and Tukey Test for TA zone (positions 2-15) are listed in Supplemental Fig. S6C and correspond to rows 40-388 in Supplemental Table S1.

### Detailed Instructions for MatrixScreener Template File

Here we detail the settings specific to our microscope and software, but these principles can be applied to other imaging modalities as well. See Supplementary Materials for a scanning template file that can be used to run an experiment using Leica LAS AF.

After LAS AF has loaded and the appropriate laser channels have been turned on, select Configuration > Settings and make sure that a) the highest *Bit Depth* is selected under *Resolution* (to offer the greatest possible dynamic range); b) *Enable During Acquisition of Series* is selected under *Manual Microscope Control* (to allow on-the-fly manual adjustment to the focus plane (Supplemental Fig. S8).

Next, from the drop-down menu on the top left corner of the screen, choose *MatrixScreener* (it should be set to *TCS SPE* by default) (Supplemental Fig. S9).

From there, navigate to the *Setup Experiment* tab in the bottom left corner and click the folder icon to load a template file (Supplemental Fig. S10). A pop-up window will confirm that MatrixScreener should load the command—click *Yes*.

Once loaded, navigate to the *Setup Jobs* tab. Here you will find the *collecting pattern* that MatrixScreener will carry out at each time point. The collecting pattern consists of four “jobs”. *Autofocus* (Supplemental Fig. S11) finds the plane with the highest contrast (using Contrast Based Method 1) and sets this as the Focus Plane that the Tracking and Acquisition jobs will reference. Its parameters are those used by Drift Correction (under the *Setup Experiments* tab). *Tracking* then takes an image of this plane, following user-specified parameters (Supplemental Fig. S12). During each loop, the Tracking job detects and draws a red box around the object to be tracked and centers this object. *Acquisition* finally images the root (Supplemental Fig. S13). The *Dud* job is required for Tracking and Drift Correction to function properly (for reasons beyond our understanding) (Supplemental Fig. S14). Given that its settings are the last used in each acquisition loop, they are copied from the autofocus job in order to minimize adjustment time (e.g. changing lasers or acquisition parameters).

In the Autofocus job, click the arrow next to *XY: 128 × 128* to expand this menu. Make sure that *Pinhole* is checked. Next, find the root tip using brightfield and the microscope eyepiece and center it. Once centered, click *Live* to visualize fluorescence.

Next, in Tracking, ensure that the focus plane is zeroed out by clicking *Set Plane* and choosing *Set Focus Plane* if that option is available (if the focus plane is already set it will display *Rest Focus Plane* instead) (Supplemental Fig. S15).

To avoid laser overexposure, the Autofocus and Tracking steps must be carried out as rapidly as possible. Given the small fluorescent footprint of QC-expressed p*WOX5::GFP(ER)*, the Autofocus range can be set to 20μM, with 20 steps (1 μM each). The pixel dimensions (“Format”) for Autofocus should be as small as possible (128 × 128) since with high Gain [V] / low Offset [%] the heterogeneities in brightness can be smoothed out such that a discrete “blob” is detected by the software. However, Tracking requires slightly higher resolution (256 × 256) so that a minimum number of pixels are present for detection. Using the Accu function to layer the intensity from several scans, the image can be multiplied to amplify the overall brightness while keeping laser power low.

The lasers are set to Seq. (sequential mode) since alternating between the two seemed to result in less photobleaching and for any given frame, and the two channels are still internally in register.

Lastly, select the *Setup Experiment* tab from which this template was loaded. On the right menu, near the top, you may enter the *Time Settings* for your particular experiment (Supplemental Fig. S16). In this example the *Total run time* is one week (168 h) and the acquisition interval (*Repeat all*) is 13m. To determine the shortest possible acquisition interval for your given experimental conditions, run a test acquisition by navigating to *Setup Jobs* (see below) and clicking *Start Record*. This will carry out an acquisition job only (without cycling through the autofocus, tracking, and dud jobs). Note the total time here and add 10 minutes to it to determine the acquisition interval (e.g. in this template the acquisition takes roughly 3m to run, with a 10 minute “cool-down” to prevent photo-bleaching and -toxicity).

Under *Start Co-ordinates* click *Learn* to center the objective on the object to be tracked. Finally, click the play icon to begin the experiment.

## Acknowledgments

The authors thank Zoé Joly-Lopez and members of the Birnbaum lab for comments on the manuscript. They also thank Scott MacCleery at Leica Microsystems for troubleshooting the imaging system.

---

**Supplemental Figure S1.** Photo of a root inside the coverslip setup described in schematic form in Figure 2. A corresponds to main Figure 2b. B corresponds to main Figure 2F. C corresponds to main Figure 2G.

**Supplemental Figure S2.** A root that had coiled in on itself having grown horizontally in the growth chamber described in Figure 1.

**Supplemental Figure S3.** Closeup of a root showing growth alongside a fishing line “guide”, which greatly minimizes rotation and coiling.

**Supplemental Figure S4.** Plot of Corrected Total Cell Fluorescence over time for each of the four films over six time points. Table (right) contains the underlying data. Blue line represents loess fit.

**Supplemental Figure S5.** Plot showing numbers of mitotic cells observed at a median section over time for each of the four films. Table (right) contains the underlying data. Blue line represents loess fit.

**Supplemental Figure S6.** Results from two-way ANOVA and Tukey Test for all tissues (A), initials + TA zone (B), and TA zone alone (C).

**Supplemental Figure S7.** Boxplots showing each of the tissue-specific division patterns for the TA zone (positions 2-15).

**Supplemental Movie S1.** Maximum projection time lapse of a root growing for one week inside the growth setup described in Figure 1. The QC is marked by *pWOX5::GFP(ER)* expression (cyan), and nuclei are visualized with *35S::H2B-mRFP1* expression (red). Supplementary Figure 1 shows a brightfield image of a similarly coiled root.

**Supplemental Movie S2.** Maximum projection time lapse of a root growing for several days, stabilized by sliding against a fishing line. Supplementary Figure 2 shows a brightfield image of a similarly stabilized root.

**Supplemental Movie S3.** Time lapse movie following an endodermal transit amplifying cell at position 3 (arrowhead), over the course of several divisions. The QC is marked by *pWOX5::GFP(ER)* expression (cyan), and nuclei are visualized with *35S::H2B-mRFP1* expression (red).

**Supplemental Movie S4.** Time lapse movie showing two sequential divisions of a cortical initial over the span of several days. The QC is marked by *pWOX5::GFP(ER)* expression (cyan), and nuclei are visualized with *35S::H2B-mRFP1* expression (red). <u>endo</u>: endodermis, <u>cort</u>: cortex, <u>epi</u>: epidermis, <u>LRC</u>: lateral root cap.

**Supplemental Movie S5.** Time lapse movie tracking a QC cell from Supplementary Movie 1 after it divides. The daughter of this QC cell is displaced into the Cortex/Endodermal Initial (CEI) position. The QC is marked by *pWOX5::GFP(ER)* expression (cyan), and nuclei are visualized with *35S::H2B-mRFP1* expression (red). Interestingly, *WOX5* expression seems diminished in both the divided QC cell and its CEI-positioned daughter, as compared to neighboring QC cells. <u>endo</u>: endodermis, <u>cort</u>: cortex, <u>epi</u>: epidermis.

